# Muscle stem cells produce a protective Fibrillin-1 matrix to prevent precocious activation

**DOI:** 10.1101/2025.08.02.666604

**Authors:** Eleni Chrysostomou, Oussama Smail, Artemis Vlachopoulou, Takuto Hayashi, Nicolas Tcheng, D. P. Reinhardt, Nicol Voermans, Frederic Relaix, Philippos Mourikis

## Abstract

Multiple biological mechanisms have been uncovered to regulate muscle stem cell quiescence, including inhibition of differentiation, adhesion-dependent anchoring, and translational control, which can be broadly classified as intrinsic or extrinsic properties. Here, we identify the matrix glycoprotein Fibrillin-1 (FBN1) as a Notch-regulated, cell-autonomous effector, essential for maintaining quiescence in muscle stem cells. Known for its causal role in Marfan syndrome (MFS), a connective tissue disorder that also presents with skeletal muscle atrophy, our work positions FBN1 as a critical niche component that protects stem cells from aberrant growth factor signalling. We demonstrate that targeted deletion of *Fbn1* in muscle stem cells leads to dose-dependent quiescence defects, characterized by loss of cellular projections, depletion of the stem cell pool, and progressive decline in muscle function. Consistently, human MFS muscle biopsies show abnormally activated satellite cells, implicating stem cell imbalance in the development of MFS-associated myopathy. Mechanistically, the loss of FBN1 upregulates TGFβ signalling, and pharmacological inhibition of this pathway using the Angiotensin Receptor blocker losartan restores the cellular and physiological defects of mutant muscles. These findings reveal a new quiescence-preserving mechanism through extracellular matrix-mediated shielding from mitogenic signals, and position stem cell dysfunction as a driver of MFS myopathy.

**Graphical abstract:** 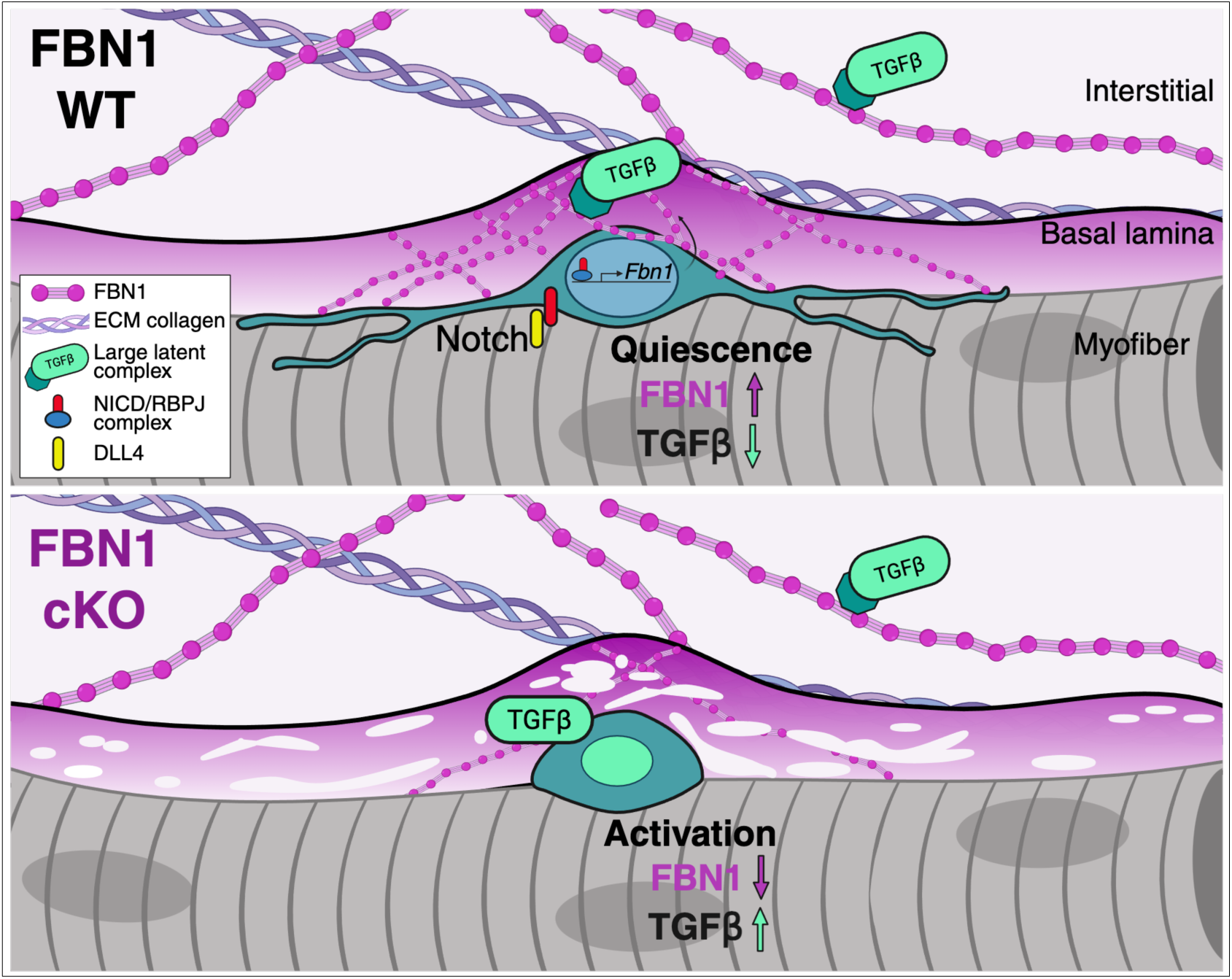

*Model of Fibrillin-1 protective barrier in quiescent satellite cells.:* Satellite cells produce Fibrillin-1 (FBN1) to establish a localized extracellular matrix barrier that limits exposure to mitogenic signals. Mechanistically, FBN1 expression is induced by Notch signalling and serves to sequester latent TGFβ complexes in an inactive form, thereby preventing their activation within the immediate niche. This barrier is spatially restricted beneath the basal lamina and remains functionally independent from the abundant interstitial FBN1 produced by fibro-adipogenic progenitors (FAPs). The model describes a self-contained, satellite cell-derived ECM system that maintains quiescence by insulating the cells from activating cues.

## Introduction

The niche of muscle stem cells, also known as satellite cells, provides both structural and signalling cues that preserve stem cell quiescence, a reversible cell cycle arrest crucial for maintaining long-term regenerative potential (Chrysostomou and Mourikis, 2024). Among the core regulators of quiescence is the Notch signaling pathway, which globally operates by suppressing activation/differentiation programs and preserving stemness. In skeletal muscle, Notch exerts its function through multiple effectors: it blocks myogenic gene expression via the *Hes*/*Hey* bHLH repressors, regulates microRNAs that ensure adhesion and anchorage, and induces the expression of extracellular matrix collagen type V, which acts as a MuSC-derived ligand for the Calcitonin receptor (Baghdadi et al., 2018a; Baghdadi et al., 2018b; Gioftsidi et al., 2022; Mourikis et al., 2012). Here, we extend this framework by identifying Fibrillin-1 (FBN1), a ubiquitous ECM glycoprotein known for its structural role and regulatory influence on growth factor signaling, as a direct target of the Notch pathway.

Mutations in FBN1 underlie Marfan syndrome (MFS), an autosomal dominant connective tissue disorder with a reported incidence of 1 in 3000 to 5000 individuals (Dietz et al., 1991; Irim Salik and Prashanth Rawla, 2023). These mutations are broadly classified as haploinsufficiency, which reduce functional protein levels or dominant-negative, where mutant FBN1 incorporates into microfibrils and interferes with their function. The variant type, however, does not reliably predict disease severity and should be interpreted with caution in clinical counseling (Arnaud et al., 2017; Milewicz et al., 2021). These mutations act either hypomorphic or dominant-negative and lead to defective mechanosensing of the cellular microenvironment and to elevated TGFβ signaling, a hallmark of MFS pathology (Milewicz et al., 2017; Neptune et al., 2003). Fbn1-deficient mice (Fbn1^−/−^) die within the first 2 weeks of postnatal life due to ruptured aortic aneurysm, impaired pulmonary function, and/or diaphragmatic collapse (Carta et al., 2006). In skeletal muscle, mice carrying one *Fbn1^C1041G^* allele, a missense mutation modeled after a mutation commonly found in MFS patients, exhibit impaired tissue regeneration (Cohn et al., 2007).

Pharmacological inhibition of TGFβ signaling has been widely explored as a therapeutic strategy in MFS and other myopathies. Among these, losartan potassium, an FDA-approved Angiotensin II receptor type 1 (AT1) antagonist, has emerged as a particularly effective candidate. Originally developed as an antihypertensive, losartan was shown to reduce TGFβ signaling, thereby indirectly antagonizing downstream fibrotic and inflammatory responses (Lim et al., 2001). In MFS mouse models, losartan prevents aortic root dilation, delays aneurysm progression, and improves cardiac, skeletal, and pulmonary phenotypes, making it one of the most extensively studied non-surgical therapies for the disease (Cohn et al., 2007; Habashi et al., 2006). In skeletal muscle, losartan enhances tissue regeneration in both Fbn1-mutant and Dystrophin-deficient (*mdx*) mice, where it mitigates fibrosis, preserves myofiber integrity, and improves muscle function (Cohn et al., 2007). To date, these effects are attributed to its capacity to create a more permissive environment for muscle stem cell-mediated repair.

Despite these results and the clear evidence of muscle atrophy in MFS patients, the direct role of FBN1 on satellite cell maintenance had not been previously investigated. Our study fills this gap, uncovering a Notch-driven, cell-autonomous function for Fibrillin-1 in maintaining stem cell quiescence and protecting against aberrant growth factor signaling. We further demonstrate that losartan not only rescues stem cell numbers and quiescence morphology in *Fbn1* cKO mice but also restores muscle performance and tissue integrity, suggesting stem cell dysfunction as a primary source of the pathology. These findings position losartan as a promising therapy for MFS-associated myopathy and provide a mechanistic basis for its beneficial effects beyond vascular tissues.

## RESULTS

### Identification of Fibrillin-1 as a potential regulator of quiescence in skeletal muscle stem cells

Notch signalling is essential for satellite cell maintenance, acting through a diverse range of targets (Gioftsidi et al., 2022). To identify additional Notch-regulated and ECM-associated regulators of satellite cell quiescence, we mined ChIP-seq data for intracellular Notch1 (NICD) and its downstream effector RBPJ in myogenic cells and integrated them with transcriptomic profiles of *in situ fixed* quiescent satellite cells (Castel et al., 2013; Machado et al., 2017). Among the genes showing both enhancer occupancy by NICD/RBPJ and strong expression in quiescent satellite cells, we identified *Fbn1*, a key extracellular matrix glycoprotein (Fig. 1A; Fig. S1A).

**Figure 1:**
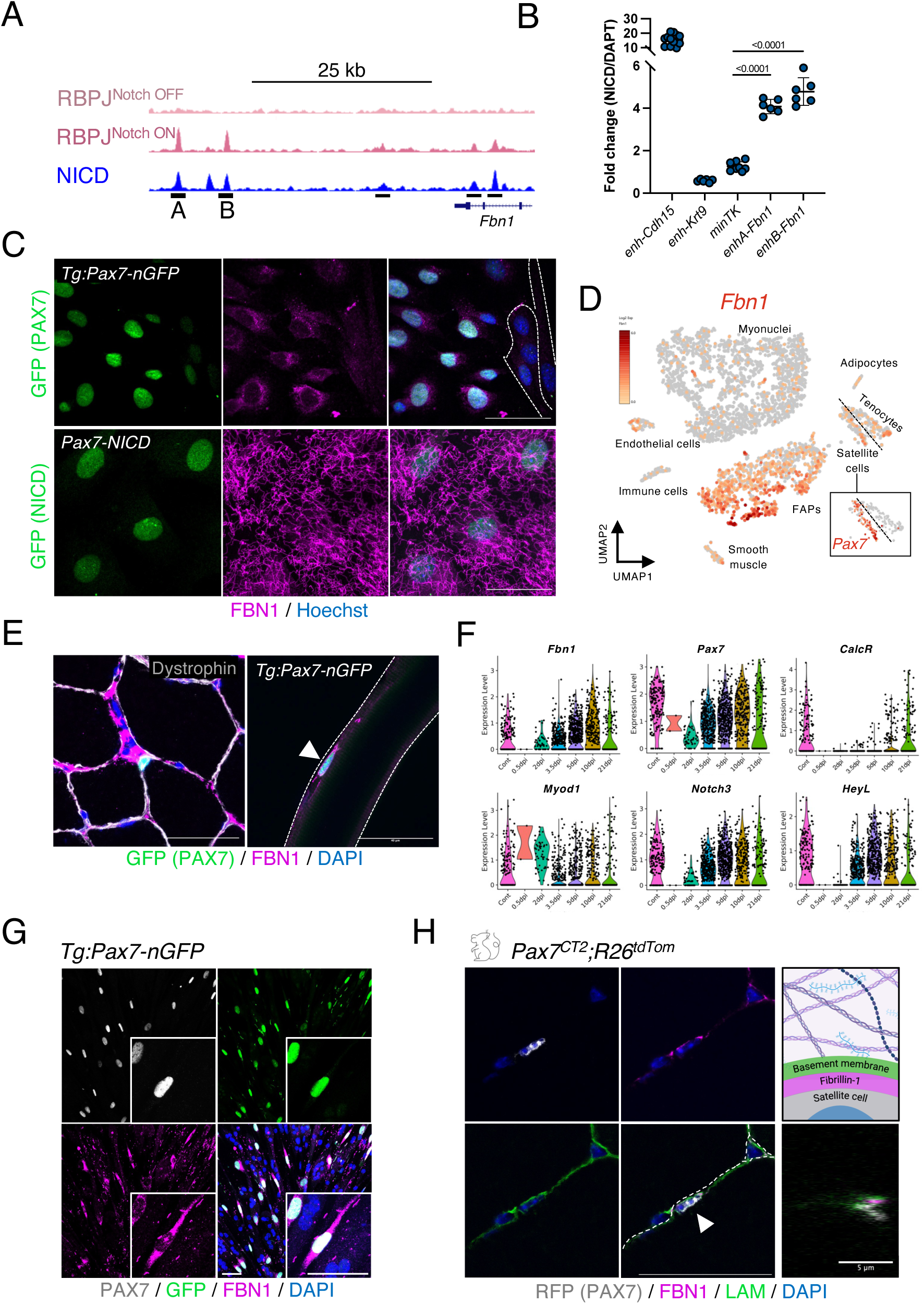
Fibrillin-1 is a Notch-regulated ECM component of quiescent satellite cells. **A.** ChIP-seq for RBPJ and nuclear Notch1 (NICD) on mouse myogenic C2C12 cells indicating enhancers associated with *Fbn1* locus (black rectangles); A and B enhancers were used for luciferase assays. **B.** Luciferase assay on cloned enhancers in C2C12 cells transfected with NICD (Notch On) or treated with the γ-secretase inhibitor DAPT (Notch Off). Cadherin 15 (Cdh15) enhancer used as positive control, Keratin 10 (Krt19) enhancer as negative control, and minTK promoter as baseline control. **C.** Isolated and cultured satellite cells from control (*Tg:Pax7-nGFP*) and Notch-overexpressing (*Pax7^CreERT2^; R26^stop-NICD^*) mice (n=3 per genotype). Dotted line indicates differentiated PAX7^-^/FBN1^-^ cells. **D.** UMAP of single-nucleus RNAseq from TA muscles (Machado et al., 2021) indicating expression of *Fbn1*. Inset shows *Pax7* expression in the satellite cell cluster **E.** Immunofluorescence for FBN1 expression in the interstitial space (TA muscle sections, left panel), and in satellite cells of isolated myofibers (100% GFP^+^/FBN1^+^, Tg:Pax7-nGFP mice, *Extensor digitorum longus* muscle; n=10-15 fibers/mouse, 3 mice). **F.** Transcriptomic data (scRNA-seq; Oprescu et al., 2020) on resting (Control) and regenerating muscles (0.5, 2, 3.5, 5, 10, and 21 days post-injury) showing the expression dynamics of *Fbn1* relative to *Pax7* (quiescence and proliferation marker), *CalcR* (quiescence marker), *Myod1* (activation and proliferation marker), and Notch pathway members *Notch3* and *HeyL*. **G.** High FBN1 expression in quiescent-like “reserve” cells (PAX7^+^) generated from FACS-isolated satellite cells (n=3 mice). **H.** High resolution imaging of TA muscle sections of FBN1 in relation to the basal lamina (LAM). Orthogonal view and schematic representation on the right-hand side. Arrowhead points to a satellite cell. Unpaired One-way ANOVA ±S.D.; scale bars = 40 μm.

Two putative enhancer elements of *Fbn1* were cloned and showed a 4- to 6-fold increase in luciferase assays upon Notch stimulation (Fig.1B). Moreover, when we isolated and cultured satellite cells from mice conditionally overexpressing NICD in satellite cells (Pax7-NICD for *Pax7^CreERT2^; R26^stop-NICD^*), we observed a striking induction of extracellular FBN1 compared to control cells (Fig. 1C). To assess the functional role of this protein, we knocked down *Fbn1* using siRNA in quiescent-like cells generated in cell culture. For that, Pax7-NICD satellite cells were isolated and cultured to confluency; sustained Notch activation prevents differentiation and fusion, leading instead to contact inhibition and subsequent cell cycle exit (Pax7⁺/EdU^-^ cells). Upon *Fbn1* knockdown, however, cells failed to maintain quiescence and prematurely entered the cell cycle, as evidenced by an increased proportion of PAX7⁺/EdU⁺ cells (Fig.S1B, C).

Single-nucleus RNA-seq data revealed that *Fbn1* is expressed by multiple cell types in skeletal muscle, with the highest levels detected in mesenchymal fibro-adipogenic progenitors (FAPs) and robust expression also observed in quiescent satellite cells (Fig. 1D). To confirm expression at the protein level, we performed immunofluorescence analysis in *Tibialis Anterior* (TA) hindlimb muscle sections using a specific antibody raised against mouse FBN1 (Shi et al., 2021). FBN1 was detected in the interstitial space, consistent with expression by FAPs, and showed complete co-localization with GFP in satellite cells on isolated myofibers from *Tg:Pax7-nGFP* mice (Fig. 1E). To assess regulation of *Fbn1* in satellite cells, we analyzed bulk RNA-seq data from Machado et al. (2017), which showed a significant reduction in expression following quiescence exit, with a 2.5-fold decrease after 120 minutes of activation (*p* = 5×10⁻⁷). This dynamic expression was further detected in single-cell RNA-seq data from injured TA muscles, where *Fbn1* expression closely correlated with Notch signaling activity, including expression of *Notch3*, *HeyL*, and the quiescence marker *CalcR* (Fig. 1F; (Oprescu et al., 2020). Moreover, FBN1 protein correlated with PAX7 in cultured cells, while expression was absent in differentiated myonuclei (Fig. 1G). Finally, to assess the regionalization of FBN1 in relation to the satellite cell niche *in vivo*, we used high resolution coupled with Single Molecule Detection confocal microscopy to visualize FBN1- and Laminin-stained sections of TA muscles and found that FBN1 deposition is polarized in the intermediate niche, located between the basement membrane and the satellite cell (Fig. 1H).

Taken together, these data indicate that Notch signaling-driven FBN1 is produced by quiescent satellite cells and deposited at the interphase of satellite cells and the basement membrane. These features suggest that FBN1 may play a cell-autonomous role in maintaining satellite cell quiescence.

### FBN1 is a functional component of the quiescent niche

To determine whether FBN1 produced by satellite cells is a functional component of the niche, we generated mice with a conditional deletion of one or both *Fbn1* alleles upon tamoxifen administration in the Pax7^+^ satellite cells, while simultaneously genetically marking them with the red fluorescence protein tdTomato (*Pax7^CreERT2^;Fbn1^+/+^;R26^tdTom^; Pax7^CreERT2^;Fbn1^fl/+^;R26^tdTom^; Pax7^CreERT2^;Fbn1^fl/fl^;R26^tdTom^*, hereafter referred to as Control, cKO^Het^, and cKO^Homo^ mice, respectively). The fidelity of the mouse model was validated by scoring Pax7^+^/Tom^+^ cells (95 %; Fig. S2A), and the reduction of *Fbn1* levels by RT-qPCR (82.5 % reduction; Fig. S2B).

Short-term chase after deletion of *Fbn1* in satellite cells (cKO^Homo^ mice, two weeks post tamoxifen treatment) resulted in their significant decrease in homeostatic muscle (47.7 % reduction), demonstrating that continuous production of FBN1 by satellite cells is essential for quiescence maintenance (Fig. 2A-C). The concomitant increase of Tom^+^ myofibers indicated that the *Fbn1* cKO cells underwent terminal differentiation and fused prematurely (Fig. 2D). Increased red fibers, but not significantly reduced satellite cells, were also scored at this time point in the cKO^Het^ mice (Fig. 2C, D). When FBN1 was depleted for a longer period (4 months post tamoxifen diet; Fig. 2A lower panel), the stem cell pool in cKO^Homo^ mice was almost completely exhausted (83.8 % reduction; Fig. 2C). Interestingly, the *Fbn1* heterozygous mice also showed a significant reduction in satellite cells, alongside with the corresponding increase in Tom^+^ myofibers (Fig. 2B, C). Long-term depletion of *Fbn1* in satellite cells also resulted in compromised architecture of the interstitial space (Fig. 2E). These global muscle defects correlated with reduced performance at treadmill endurance and grip strength tests (Fig. S2C, D). Finally, no statistically significant differences were observed at the fiber cross-sectional area (CSA) in both short- and long-term *Fbn1* depletion cohorts, although cKO^Homo^ mice showed a tendency to have larger myofibers (Fig. S2E, F).

**Figure 2:**
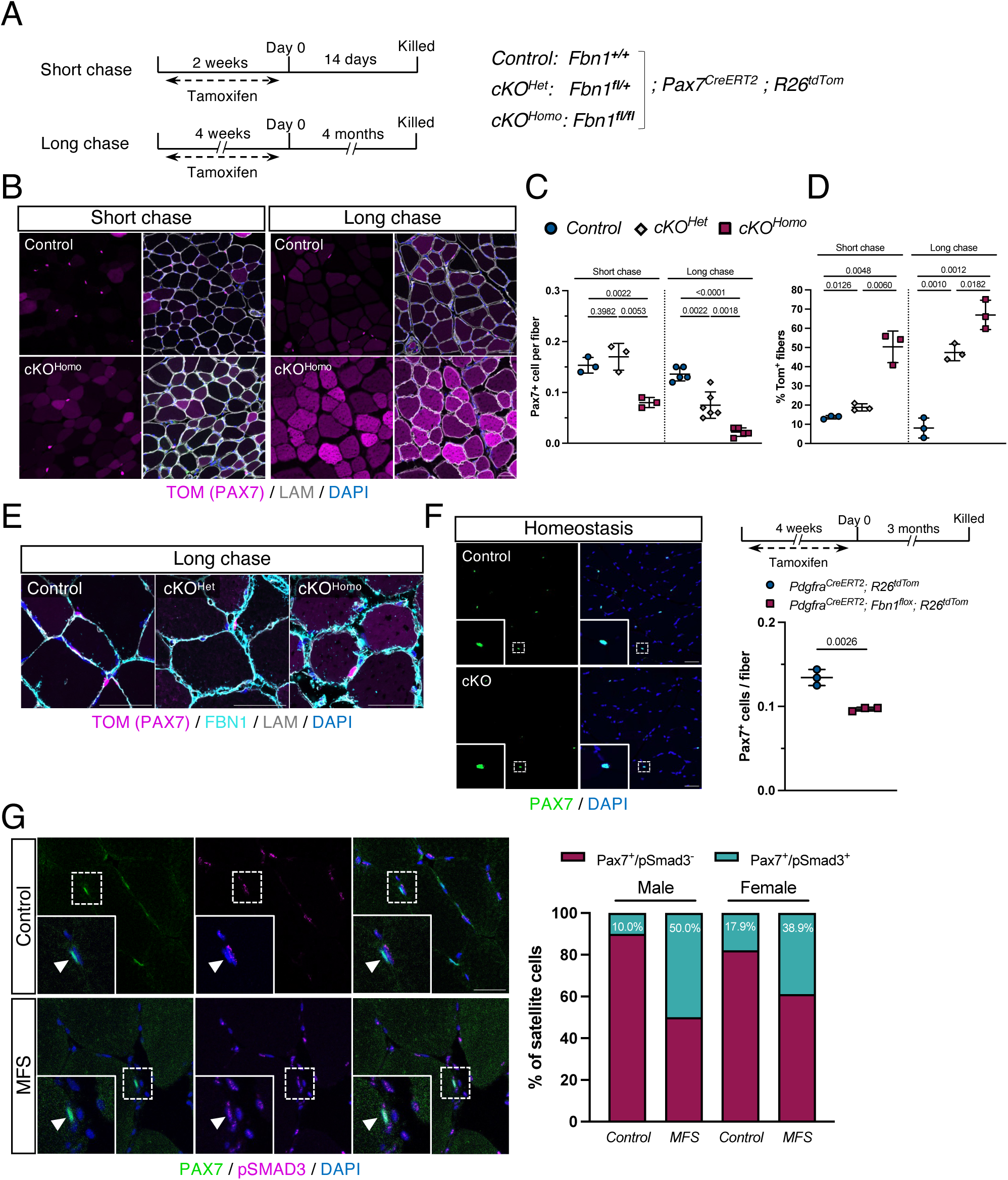
Satellite cell-derived FBN1 maintains quiescence and prevents premature fusion *in vivo*. **A.** Experimental scheme for short (2 weeks post tamoxifen treatment) and long (4 months post tamoxifen) chase in control (*Pax7^CreERT2^;Fbn1^+/+^;R26^tdTomato^*), *Fbn1* heterozygous (cKO^Het^: *Pax7^CreERT2^;Fbn1^fl/+^;R26^tdTomato^*) and *Fbn1* homozygous conditional knockout (cKO^Homo^: *Pax7^CreERT2^;Fbn1^fl/fl^;R26^tdTomato^*) satellite cells. **B.** Immunostaining of *Tibialis anterior* (TA) muscle sections from Control and Fbn1 cKO^Homo^ mice from short and long chase after *Fbn1* deletion (n = 3 mice/genotype). **C.** Satellite cell quantification in homeostatic TA muscles in a Fbn1 allelic series (n = 3-6 mice per genotype). **D.** Quantification of Tom^+^ fibers from homeostatic TA muscles in short- and long-chased animals (n = 3 mice/genotype). **E.** Immunostaining of TA muscle sections from *Fbn1* long-chase depletion mice (n = 3 mice/genotype). **F.** Immunostaining of TA muscles from control (*Pdgfra^CreERT2^;Fbn1^+/+^;R26^tdTomato^*) and *Fbn1* cKO in FAPs (*Pdgfra^CreERT2^;Fbn1^fl/fl^;R26^tdTomato^*) mice from long chase after *Fbn1* deletion (n = 3 mice/genotype), and satellite cell quantification. **G.** Immunostaining of *Vastus lateralis* muscles from age- and sex-matched control and MFS patients (arrowhead indicates a satellite cell); (*right*) quantification of PAX7^+^/pSMAD3^±^ satellite cells. One-way ANOVA ±S.D (C, D); unpaired *t-test* ±S.D (F); scale bars = 40 μm.

In the adult mouse, *Pax7* is expressed in tissues beyond skeletal muscle, including a subset of undifferentiated spermatogonia, the central nervous system and in the pituitary gland, a key paracrine regulator (Aloisio et al., 2014; Stoykova and Gruss, 1994). For that, we considered the possibility that systemic effects could arise from inadvertently targeting *Fbn1* in *Pax7*-expressing cells outside the muscle. To avoid misinterpreting the muscle phenotypes due to such paracrine effects in the *Pax7^CreERT2^; Fbn1^flox^* mice, we examined the pituitary gland where *Fbn1* is expressed. Based on published single-cell RNAseq data (Mayran et al., 2019), *Pax7* and *Fbn1* are not co-expressed in the same cell populations (Fig. S2G). To validate this in our model, we isolated the pituitary gland and performed immunostaining on sections for both PAX7 and FBN1. Consistent with the transcriptomic data, we observed no co-expression of the two proteins at the cellular level (Fig. S2H). The involvement of testicular PAX7^+^ cells was excluded since we did not observe a sex bias.

To further investigate the influence of systemic *Fbn1* depletion on satellite cells, we selectively targeted *Fbn1^flox^* in mesenchymal cells. To achieve FBN1 knockdown, we generated compound mice carrying the *Pdgfra^CreERT2^* driver and the *Fbn1^flox^* allele (Fig. S2I). Three months after tamoxifen administration, we observed a significant reduction in the number of PAX7⁺ cells in the cKO muscles (Fig. 2F). It is important to note that *Pdgfra* is broadly expressed across multiple tissues, and *Pdgfra-Fbn1* cKO mice developed pathological phenotypes (like abdominal hernias; data not shown). Therefore, the observed decrease in satellite cell abundance could result from *Fbn1* loss specifically in FAPs but may also reflect broader systemic dysregulation or circulating signals triggered by widespread FBN1 deficiency.

We also examined the quiescent status of satellite cells in human biopsies from *Vastus lateralis* muscles from healthy individuals and MFS patients. Biopsies were obtained from two MFS patients, a 58-year-old female and a 32-year-old male, as well as from age- and sex-matched healthy controls, a 54-year-old female and a 31-year-old male. The MFS patients carried missense mutations in FBN1, located in exons 6 and 63 respectively (Voermans et al., 2009). Consistent with our findings in mice, satellite cells from MFS patients exhibited elevated levels of nuclear phosphorylated SMAD3 (pSMAD3) (Fig. 2G). Although both MFS samples showed increased pSMAD3 staining, the number of satellite cells appeared unchanged relative to controls (Fig. S2J). Notably, *Fbn1^+/–^* mice also showed increased pSMAD3 in satellite cells, yet this was accompanied by a progressive, age-related decline in satellite cell number. One possible explanation for this discrepancy lies in the nature of the mutation, as the mouse model carries a null allele of *Fbn1*, whereas the MFS patients harbor heterozygous missense mutations, which may have different effects in satellite cells.

### Loss of FBN1 leads to TGFβ activation of satellite cells

It has been shown that FBN1 can sequester latent TGFβ complexes in the ECM, and upon its destabilization, bioactive TGFβ is released from the ECM (Annes et al., 2003). Whether a similar mechanism operates in satellite cells remained unknown. Using pSMAD3 as a reporter of TGFβ signalling activity, we found cKO satellite cells with ectopic pathway activation. This response was FBN1 dose-dependent with cKO^Het^ muscles exhibiting almost half the number of pSMAD3^+^ PAX7^+^ cells compared to cKO^Homo^ (Fig. 3A, B). Surprisingly, the mutant cells despite being activated (pSMAD3^+^) and fusing (TOM^+^ fibers), bypassed the proliferation phase and directly underwent fusion while remaining Ki67⁻ (Fig. 3C), a phenotype that, to date, has only been described in *Rbpj* mutant satellite cells (Bjornson et al., 2012; Mourikis et al., 2012).

**Figure 3:**
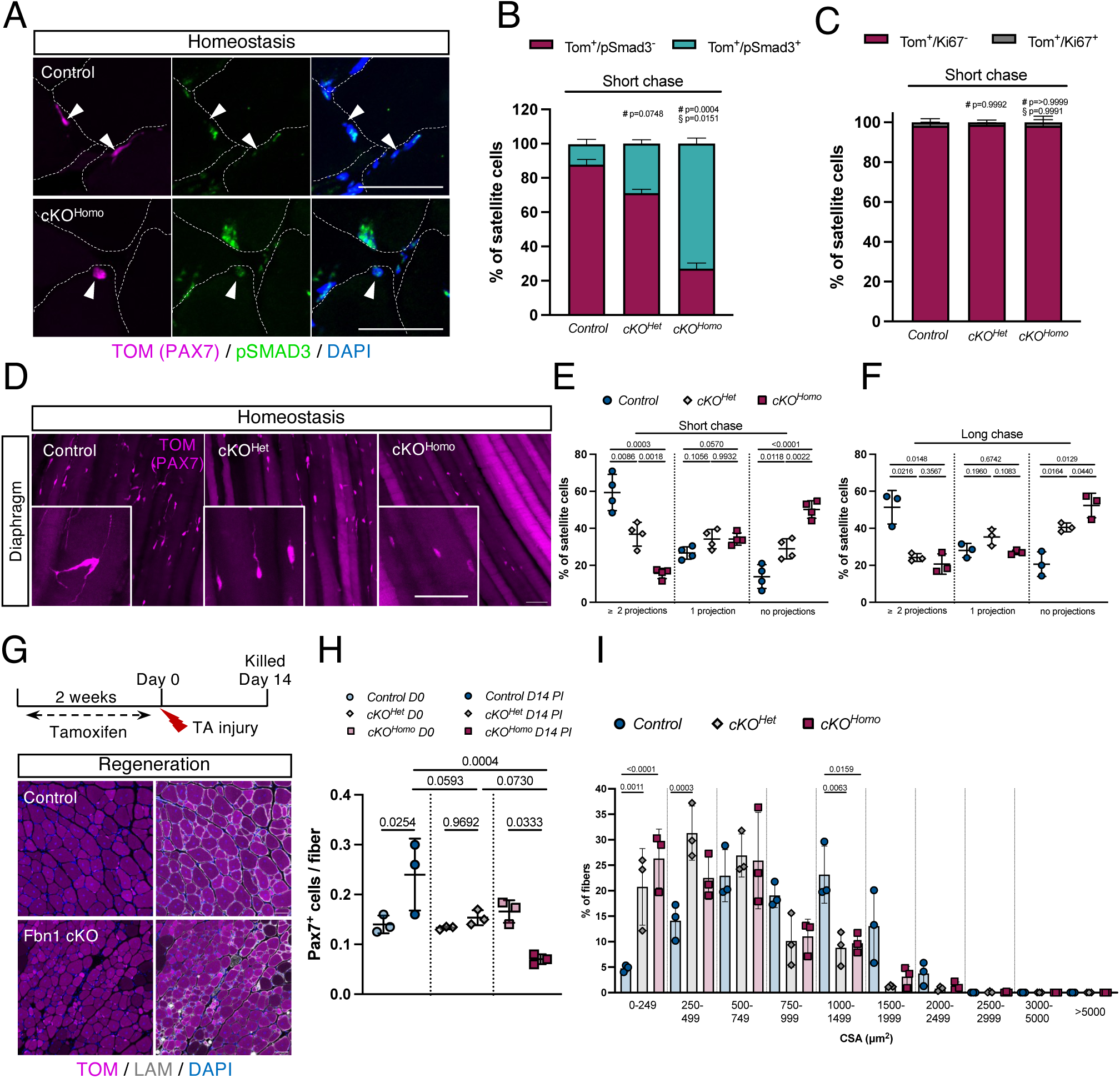
Loss of FBN1 activates TGFβ signaling in satellite cells and disrupts quiescence projections. **A.** Immunostaining of resting TA muscle sections from Control and *Fbn1* cKO^Homo^ TA 2 weeks after tamoxifen-induced *Fbn1* deletion. Dotted lines outline the myofibers. Arrowheads point TOM^+^ satellite cells. **B.** Quantification of Tom^+^/pSmad3^±^, and **C.** Tom^+^/Ki67^±^ satellite cells in Control and *Fbn1* mutant satellite cells (n = 3 mice/genotype). **D.** Immunostaining of diaphragm muscles from control and cKO Fbn1 mice (short-term chase). **E.** Quantification of satellite cell projections divided in three groups: ≥2 projections, 1 projection, no projections, in short (2 weeks) and **F.** in long (4 months) chase after *Fbn1* deletion. **G.** Experimental scheme of muscle injury with BaCl_2_ and 14 days of regeneration. **H.** Quantification of PAX7^+^ cells in TA muscle sections at 0 and 14 days post injury (n = 3 mice/genotype). **I.** Quantification of fiber cross-sectional area (CSA) from the short-term, regenerative Fbn1 depletion cohort (n = 3 mice per genotype). One-way ANOVA ±S.D.; # compared to Control; § compared to cKO^Het^; scale bars = 40μm.

Given this atypical activation state, we scored satellite cells for cytoplasmic projections, a hallmark of an early activation response (Kann et al., 2022; Ma et al., 2022). For that, we examined the diaphragm, a thin, easily accessible muscle well-suited for whole-mount imaging and rich in PAX7^+^ cells. As shown in Figure 3D-F, most *Fbn1* cKO satellite cells had exited quiescence, as indicated by the attenuation of the quiescent projections. This response was also strictly dose-dependent, with cKO^Het^ satellite cells exhibiting an intermediate phenotype between control and cKO^Homo^ cells. Moreover, the projections defect in Fbn1 heterozygous cells gradually aggravated with time, confirming Fbn1’s haploinsufficiency in the mouse.

To investigate the regeneration and renewal potential of *Fbn1* cKO satellite cells, we performed BaCl_2_-mediated injuries of TA muscles and analysed them 14 days later (Fig. 3G). Despite abundant FBN1 in regenerating muscle likely produced by resident fibroblasts (Fig. S3A), fewer self-renewing PAX7^+^ cells were observed in the *Fbn1* cKO mice (Fig. 3G, H), confirming a cell-autonomous role for FBN1. In addition, *Fbn1* cKO satellite cells failed to return to quiescence during regeneration as indicated by the increased levels of pSMAD3 and, to our surprise, the proliferation levels remained the same between control and cKO cells (Fig. S3B, C). Notably, mutant cells produced significantly smaller nascent myofibres compared to control cells (Fig. 3I). To investigate self-renewal in a more tractable and fibroblast-free system, we targeted *Fbn1* in satellite cells using short-interfering RNA on isolated myofibres in culture. Consistent with our *in vivo* observations, *Fbn1* knockdown resulted in a marked decrease in the number of the self-renewing PAX7^+^MYOD^−^ cells (10.5 %), compared to scramble control (27.7 %), (Fig. S3D-F).

### Angiotensin II type 1 receptor antagonist restores quiescence of Fbn1-deficient satellite cells and prevents muscle performance decline

Previous studies have shown that administration of losartan in multiple myopathic states can restore muscle architecture and prevent aortic aneurisms in MFS mouse models (Biressi et al., 2014; Elbadawi et al., 2019; Lim et al., 2001). Also, in both homeostatic and regenerating muscles, losartan has been shown to abrogate TGFβ signaling and restore muscle strength (Cohn et al., 2007). However, the mechanisms underlying these effects are not well characterized, and the existing data on satellite cell behavior may be misleading due to the use of markers that are not broadly validated for distinguishing quiescence, activation, and differentiation in the muscle field (Cohn et al., 2007).

Our genetic experiments demonstrated that targeted deletion of *Fbn1* in satellite cells leads to ectopic activation of TGFβ signaling, depletion of the stem cell pool, and severe impairment of muscle homeostasis and function. To test whether losartan could mitigate these effects, we administered the drug to both mutant and control mice using short- and long-term treatment regimens. We first validated efficient TGFβ signaling inhibition by losartan in satellite cells (Fig. S4A-C). Notably, the losartan-target receptor AT1 is not expressed in either quiescent satellite cells or activated myoblasts (Fig. S4D). This suggests that losartan’s inhibitory effect on TGFβ signaling is AT1-independent or results from systemic effects, such as a reduction in circulating TGFβ levels, as previously reported (Campistol et al., 1999; Morita et al., 2024).

Treatment of *Fbn1* mutant mice with losartan for four weeks (short-term) prevented satellite cell loss (Fig. 4A-C), while pSMAD3 in Pax7^+^ cells in both cKO^Het^ and cKO^Homo^ muscle was restored to levels comparable to control-treated mice (Fig. 4D). Induction of TGFβ signalling is linked to proliferating muscle cells (Fig. S4B, C), whereas the retraction of the quiescent projections is a hallmark of early activation, which occurs before proliferation (Girardi et al., 2021; Kann et al., 2022; Ma et al., 2022).To measure the impact of losartan on satellite cell activation state, we scored as previously the cytoplasmic projections and compared treated muscles to untreated. Losartan shifted the satellite cell population towards a quiescent state (≥2 projections) and away from an activated state (no projections) (Fig. 4E-F). No statistically significant differences were observed in the fiber cross-sectional area (CSA) between genotypes, with a tendency towards bigger myofibers in the cKO^Homo^ mice (Fig. S4E). Compared to the non-treated mice (Fig. S2E), losartan-treated myofibers fully shifted towards bigger CSA levels possibly due to the inhibitory role of TGFβ pathway on myocyte fusion (Girardi et al., 2021). Losartan did not show any obvious signs of adverse effects on satellite cells as Control and Control-losartan cells were very similar in all sets of measurements (Fig. 4C, D, F).

**Figure 4:**
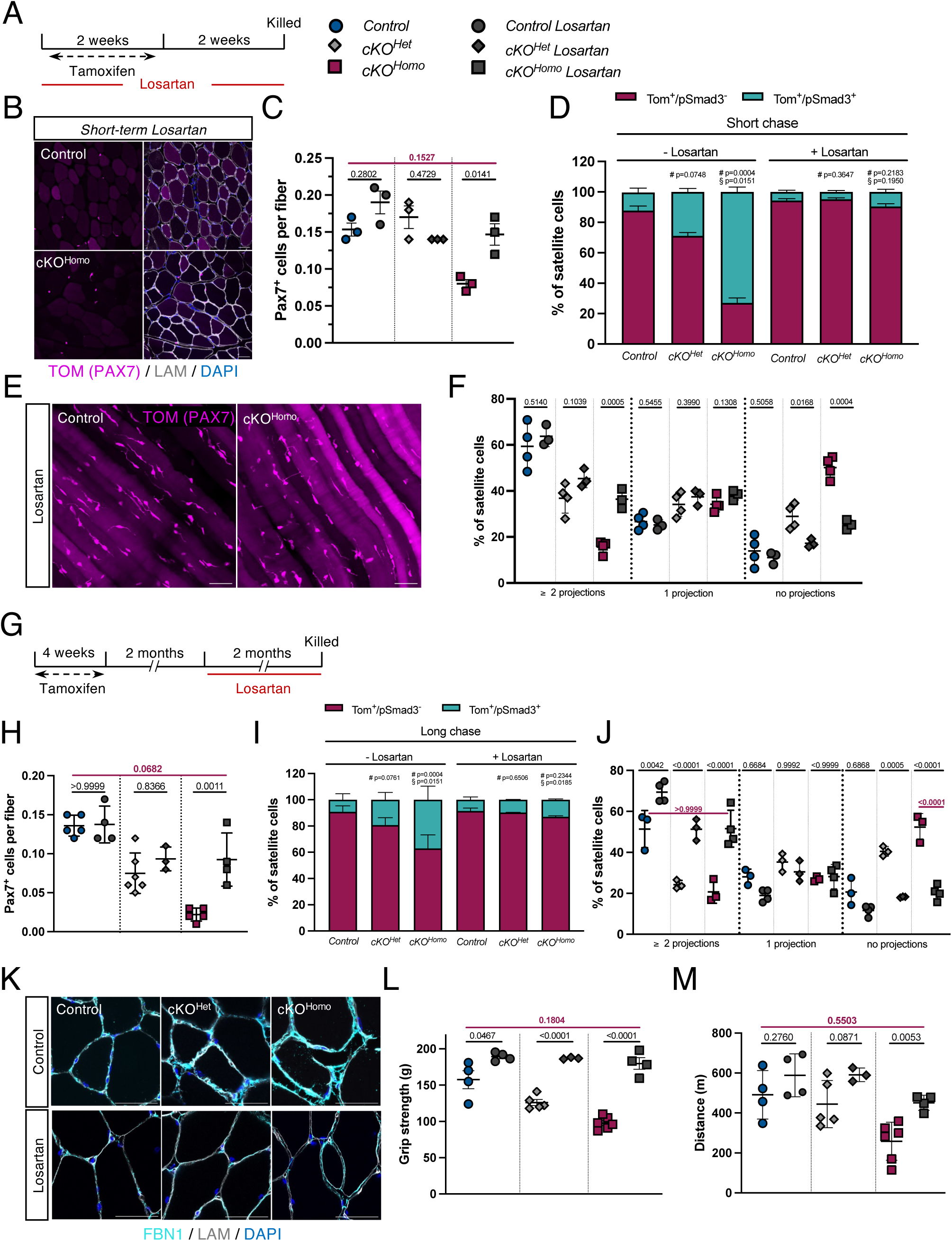
Angiotensin II type 1 receptor antagonist restores quiescence of Fbn1-deficient satellite cells and prevents muscle performance decline. **A.** Experimental schematic of short-term exposure (4 weeks) to losartan in *Fbn1* cKO mice. **B.** Immunostaining of Control and *Fbn1* cKO TA muscle sections from short-term losartan exposure (n = 3 mice/genotype). **C.** Satellite cell quantification and **D.** Tom^+^/pSmad3^±^ in homeostatic TA muscles (n = 3-6 mice/genotype) with and without exposure to losartan (n = 3 mice/genotype per treatment). **E.** Immunostaining of diaphragm muscles from Control and *Fbn1* cKO mice treated with losartan short-term, and **F.** Quantification of satellite cell projections in mice treated with losartan short-term. **G.** Experimental schematic of long-term (2 months) exposure to losartan in *Fbn1* cKO mice, starting at 2 months after end of tamoxifen treatment. **H.** Satellite cell quantification and **I.** Tom^+^/pSmad3^±^ in homeostatic TA muscles with and without exposure to losartan (n = 3-6 mice/genotype per treatment). **J.** Quantification of satellite cell projections in mice treated with losartan long-term. **K.** Immunostaining of TA muscle sections from Control and *Fbn1* cKO TA muscles from cohorts with and without losartan treatment **L.** Quantification of grip strength and **M.** treadmill performance from the long-term *Fbn1* depletion cohort on losartan (n = 3-6 mice per genotype). Losartan: 0.6 g/L in drinking water; one-way ANOVA ±S.D.; # compared to Control; § compared to cKO^Het^; Scale bars = 40 μm.

We also treated *Fbn1* mutant mice with losartan for an extended period (8 weeks), starting at 2 months after tamoxifen induction (Fig. 4G). The rationale for this timing was to assess whether losartan could restore the satellite cell population or prevent its depletion. This approach also aligned with clinical practice, as Marfan patients typically begin losartan treatment after disease onset (Elbadawi et al., 2019). The number of PAX7^+^ cells in cKO^Homo^ muscles showed a significant degree of rescue (Fig. 4H; Fig. S4F). Moreover, in both cKO^Het^ and cKO^Homo^ groups pSMAD3 levels were restored and were comparable to control groups, indicating potent inactivation of TGFβ signaling in *Fbn1* cKO satellite cells and return to quiescence levels (Fig. 4I). This was further supported by the full reestablishment of projections in the remaining satellite cells across the board (Fig. 4J).

Encouraged by the rescue results on the cellular level, we next evaluated the impact of losartan treatment on muscle architecture and functional performance. As shown in figure 4K, losartan treatment resulted in almost complete restoration of muscle morphology, and functional assays using grip strength and treadmill endurance tests demonstrated partial to full recovery of muscle performance and strength (Fig. 4L, M). By measuring the CSA of the myofibers, we did not observe any statistically significant difference between the size groups and genotypes, but a clear shit in considerable bigger myofibers was observed compared to non-treated mice (compare Fig. 4E, G to Fig. S2E, F).

## Discussion

A central challenge in stem cell biology is to uncover the diverse strategies that stem cells have evolved to preserve their regenerative potential and quiescent state within dynamic and physiologically demanding tissue environments. In satellite cells, intrinsic properties and extrinsic signals, such as transcription factors or factors secreted by neighboring myofibers, contribute to their maintenance. Most of the factors identified to date that regulate satellite cell quiescence act to reinforce quiescence, either by preventing differentiation (Notch signaling), ensuring correct anchoring (Integrinβ1 and Cadherins M/N), or selectively enabling translation of specific mRNAs required for the quiescent state (initiation factor 2 alpha eIF2α), (Goel et al., 2017; Mourikis et al., 2012; Rozo et al., 2016; Zismanov et al., 2016). Some mechanisms, however, function specifically to prevent early activation. For example, CalcR maintains satellite cell quiescence by sensing collagen V in the niche and activating Lats1/2, which phosphorylates and inhibits YAP, thus blocking the transcriptional programs required for activation (Baghdadi et al., 2018a; Zhang et al., 2019). Similarly, the RTK signaling antagonist Sprouty1 preserves satellite cell quiescence by inhibiting FGF-induced ERK signaling, thus preventing their activation (Chakkalakal et al., 2012). Our findings reveal a new paradigm in stem cell regulation and expand the panel of quiescence mechanisms regulated by Notch signalling (Gioftsidi et al., 2022). Here, we show that Notch-driven FBN1 deposition forms a sub-basal lamina “quiescence dome”, a protective extracellular matrix barrier assembled by the stem cells themselves, that insulates them from mitogenic cues in the environment (Fig. S4H). This cell-autonomous niche component complements previously known quiescence regulators and underscores the active role of the stem cells in constructing their own microenvironmental niche.

We also propose that the cell-autonomous function of FBN1 is preserved during tissue regeneration. In resting muscle, satellite cells reside tightly confined between the myofibre and its basal lamina, physically separated from the interstitial space where FBN1 is abundant. This defined anatomical niche is drastically remodeled following injury, with myoblasts, mesenchymal cells and infiltrating immune cells intermingling, and FBN1 remaining abundant or even increased throughout the tissue. Nevertheless, Pax7⁺ progenitors lacking Fbn1 fail to self-renew, demonstrating an absolute cell autonomous requirement for Fbn1 production both in quiescent and proliferating progenitors.

Notably, the accepted current model for thoracic aortic diseases such as MFS is that the primary driver is defective adhesion of smooth muscle cells to a compromised ECM, with TGFβ overactivation representing a secondary repair response (Milewicz et al., 2017). In contrast, we found that inhibition of TGFβ signaling in *Fbn1* cKO satellite cells fully restored their quiescent projections, which are adhesion-dependent structures that sense niche biomechanics (Krauss and Kann, 2023). Restoration of quiescence with robust cytoplasmic projections in satellite cells lacking FBN1 argues against a model in which TGFβ hyperactivation is merely secondary to defective adhesion. Collectively, our analyses of *Fbn1* cKO mice and their response to losartan suggest that FBN1 functions primarily as a signaling regulator rather than a mechanical element of the satellite cell niche.

Beyond the specific context of muscle, since FBN1 is widely expressed in various tissues, a pertinent question is whether similar autocrine matrix-based quiescence mechanisms operate in other stem cell compartments and developmental processes. In the developing retinal vasculature, FBN1 acts at the angiogenic front that regulates endothelial tip/stalk specification by reducing VEGF-induced Delta-like 4 induction and Notch signalling, largely through its influence on associated proteins like MAGP1 and ADAMTS1 (Alonso et al., 2023). Also, in the developing marrow niche FBN1 modulates TGFβ availability, thereby regulating skeletal stem cell activity (Smaldone et al., 2016). Notably, while systemic administration of a TGFβ-neutralizing antibody improved bone mass and restored marrow adipogenesis, treatment with losartan did not alleviate bone loss in MFS mice (Nistala et al., 2010; Ramirez et al., 2018). Instead, we found that losartan rescued the depletion of the satellite cell pool and restored both quiescence and muscle function. It would be of great interest to investigate whether the differential responses of various stem cell types reflect distinct niche properties, as well as intrinsic differences between proliferative (emerging) and non-proliferative (quiescent) tissue environments.

Finally, our findings provide new insight into the pathophysiology of Marfan syndrome myopathy. We observed that muscle biopsies from MFS patients contain activated satellite cells, indicating a compromised stem cell pool in the disease. Encouragingly, the losartan rescue of satellite cell function and muscle performance in our mouse model hints at a therapeutic opportunity.

Therefore, losartan, already in clinical use for preventing aortic aneurysm in MFS, may also alleviate muscle degeneration by normalizing the satellite cell niche signaling environment. Our work thereby positions muscle stem cell dysfunction as a previously underappreciated driver of MFS myopathy.

## Author contribution

EC conceptualised the work, performed the experiments, analysed the data and wrote the original manuscript; PM conceptualised and supervised the work, obtained funding and wrote the manuscript; F.R obtained funding and edited the manuscript; OS carried out the experiments related to mesenchymal cells and AV those related to the pituitary gland and the standardisation of losartan; TH carried out the scRNAseq data integration analysis; and DPR provided the FBN1 antibody and edited the manuscript; NV provided MFS muscle biopsies.

## Funding

This work was supported by the French National Research Agency (ANR) grants (ANR-19-CE13-0010 and ANR-19-CE14-0008), the Labex-REVIVE (ANR-10-LABX-73), the International Research Project of INSERM Calci-Notch, the Association Française contre les Myopathies (AFM) via TRANSLAMUSCLE (PROJECT22946), and the AFM-Telethon postdoctoral fellowship (28851). N.C.V. was supported by an NWO (Netherlands Organisation for Scientific Research) grant in 2006. DPR received funding from the Genetic Aortic Disorders Association Canada.

## Conflict of interest statement

The authors declare no conflict of interest.

## Acknowledgement

The authors would like to thank F. Ramirez for kindly supplying the *Fbn1^flox^* mice, the animal units of the IMRB and LEAT of the Structure Fédérative de Recherche (SFR), Necker (Paris, France), and the histology, imaging and image analysis platforms of the SFR Necker. Finally, we would like to thank Baptiste Periou and Pr F.-J. Authier for providing us with the control human muscle biopsies

## Methods

### Mouse strains

The mouse lines used in this study have been previously described and kindly provided as described: *Tg:Pax7-nGFP* (S. Tajbakhsh; (Sambasivan et al., 2009), *R26R^stop-NICD-nGFP^* (D. Melton; Jackson Laboratories, stock 008159; (Murtaugh et al., 2003)), *Pax7-^CreERT2^* (G. Kardon; Jackson Laboratories, stock 017763; (Murphy and Kardon, 2011)), *Pdgfra-^CreERT2^* (Chung et al., 2018), *R26R^td Tomato^* (Madisen et al., 2010), *Fbn1^flox^* (F. Ramirez; (Cook et al., 2012)). Animals were handled as per French and European Community guidelines and protocols were approved by the ethics committee at the French Ministry (Project No: 20-027 #24357). Mice used were 6-12 weeks old.

### Human muscle biopsies

Muscle biopsies were obtained from diagnosed MFS described in (Voermans et al., 2009)., with protocol number CMO Radboudumc number (CMO-nr.): 2005/311.

### Muscle injury

Mice were anaesthetized with isoflurane using the open-drop and nose cone method. Once the mouse was confirmed unresponsive to noxious stimuli (i.e. toe pinch), the lower part of the leg was shaved for better visualization, and the TA muscle was injected with 50μl of 1.2% BaCl_2_ (Sigma, 202738) using a 28-gauge needle.

### Tissue fixation for immunofluorescence

*Tibialis anterior* (TA) and diaphragm (DIA) muscles were fixed in 2% PFA/PBS for 2 hours at 4° C and washed 3 times with 1X PBS. DIA muscles were then directly subjected in wholemount staining (see section “Immunofluorescence staining”). TA muscles were immersed in 15% sucrose/PBS overnight, transferred in 30 % sucrose/PBS the following day for 4-5 hours, embedded in OCT, frozen in liquid-nitrogen-cooled isopentane and sectioned transversely at 8 μm. Pituitary glands were fixed in 4% PFA/PBS for 12 hours at 4°C, washed 3 times with 1X PBS, and further processed as TA and DIA. Human muscle biopsies were frozen fresh. Once thawed they were fixed in 4% PFA/PBS overnight at 4° C, washed 3 times with 1X PBS, embedded in paraffin and sectioned transversely at 4 μm.

### Single myofiber isolation and siRNA transfection

Individual myofibers were isolated from *Extensor digitorum longus* (EDL) muscles following a modified version of the previously described protocol (Shinin et al., 2006). Briefly, EDL muscles were incubated at 37°C in 0.2% collagenase type I (Sigma, C0130) in isolation medium (GlutaMAX DMEM, Thermo, 61965069) without antibiotics for 2 h. Following enzymatic digestion, individual myofibers were released by mechanical dissociation, transferred to serum-coated Petri dishes, and transfected with *Fbn1* siRNA (Dharmacon SMARTpool; ON-TARGETplus Fbn1 (14118) L-045295-01-0020) or scramble siRNA (Dharmacon ON-TARGETplus Non-targeting D-001810-01-20) at a final concentration of 200 nM, using Lipofectamine 2000 (ThermoFisher, 11668) in Opti-MEM (Gibco, 31985-070). The following day, the fibers were placed in culture medium (isolation medium with 1%penicillin/streptomycin (GIBCO), 20% FBS, 1% rat muscle extract), incubated for 72 hours at 37°C / 3% O_2_ and fixed in 4% PFA/PBS before immunostaining.

### Isolation, culture and manipulation of satellite cells

Hindlimb muscles were dissected and enzymatically digested in Dispase A and Collagenase II as previously described (Baghdadi et al., 2018). Satellite cells were isolated by FACS (85 μm nozzle) using either GFP or RFP markers depending on the mouse line. Isolated satellite cells were plated on Matrigel-coated (Corning, 354248), 8-chamber slides (Sarstedt, 94.6140.802) in growth medium composed of DMEM (GIBCO) with 20% FBS, and 1% penicillin/streptomycin (GIBCO), at 37° C. For *Pax7^CreERT2^; R26^stop-NICD-ires-nGFP^*, prior FACS, bulk muscle preparations were pre-cultured for 3 days. siRNA transfection of isolated satellite cells was performed as described in “Single myofiber isolation and siRNA transfection” section. Satellite cells from this mouse line were cultured for 6 days before siRNA. Following siRNA transfection, cells were pulsed with 10 μM EdU (Click-iT PLUS Kit C10640, Life Technologies) for 24 h before fixation. For reserve cell cultures, isolated satellite cells were cultured for 10-14 days (as described in Baghdadi et al., 2018) and fixed in 4% PFA/PBS for 10 min, followed by 3 washes in 1X PBS before immunostaining.

### Immunostaining

After fixation, cells, myofibers and diaphragms were washed three times with 1X PBS and then permeabilized and incubated in blocking solution containing 5% BSA and 0.25% Triton X-100 in PBS for 60 min in RT. Following, the samples were incubated with primary antibodies (Table S1) overnight at 4° C. After 3 washes with 1X PBS, samples were incubated with the corresponding secondary antibodies and Hoechst for 1h at RT. EdU staining was performed right after the secondary antibody incubation according to manufacturer’s instructions (Click-iT PLUS Kit C10640). Laminin staining was performed with a conjugated version (NB300-144AG647, Bio-techne) after secondary antibody incubation for 2 h at 37° C followed by Hoechst staining and mounting in fluoromount (00-4958-02, Invitrogen). Tissue sections from TA and pituitary glands were treated the same way as described above with an additional fixation before the blocking step in 4% PFA/PBS.

### Construction of luciferase reporter and luciferase assays

Candidate enhancers of *Fbn1*, *Keratin9* and *Cadherin15* were amplified by PCR from genomic DNA of C2C12 cells. The enhancers were inserted into the firefly-luciferase pGL3-Basic vector (Promega, E1751) upstream of a minimal thymidine kinase promoter and then sequenced. C2C12 cells were transfected using Lipofectamine LTX (Life Technologies, 15338030), lysed and the luciferase signal was measured using the Dual-Luciferase Reporter Assay System (Promega, E1910).

For normalization, *Renilla* luciferase (pCMV-Renilla) was transfected at 1:20 ratio relative to firefly-luciferase constructs. The sequences of enhancers are listed in Table S2.

### RNA isolation and RT-qPCR

Total RNA was extracted from satellite cells isolated by FACS using RNA/DNA Microprep kit (Zymo Research, R1051) and reverse transcribed using SuperScript III (Invitrogen, 18080093), as per the manufacturer’s instructions. RT–qPCR was performed using FastStart Universal SYBR Green Master mix (Roche, 04913914001). Primers used for qPCR analysis are listed in Table S3.

### Single-cell RNA-seq data analysis

Publicly available single-cell RNA-sequencing (scRNA-seq) data from regenerating mouse skeletal muscle were obtained from the study by (Oprescu et al., 2020). The original dataset was downloaded from the NCBI Gene Expression Omnibus (GEO) under accession number [GSE 138826]. All downstream analyses were performed using R (version 4.4.2) and the Seurat package (version 5.3.0) (Hao et al., 2024). Raw count matrices were processed following standard Seurat workflows, including quality control, normalization, dimensionality reduction, clustering, and visualization. Cell type annotation was based on the original metadata provided in the publication and further refined using canonical marker genes. *Pax7*-positive cells were first defined as satellite cells, and this population was extracted using the subset() function in Seurat. The subsetted satellite cells were then reclustered, and clusters characterized by high Pax7 expression were identified and isolated as quiescent satellite cells (qSCs). Violin plots of gene expression during muscle regeneration were generated using these qSCs, as shown in Figure 1F. All custom R scripts and analysis pipelines used in this study are available as supplementary file (Supplementary Code S1).

### Losartan treatment

Six-week-old mice were administered Losartan potassium (Sigma-Aldrich, Cat. #L1299) via drinking water at a concentration of 0.6 g/l, or received placebo (regular water). Drinking bottles were protected from light to preserve drug stability and the losartan solution was replaced twice weekly for the entire duration of the treatment. Water consumption was monitored throughout the study, and no significant differences were observed between placebo and Losartan-treated groups.

### Muscle performance tests

For the treadmill test, mice were pre-acclimatised for five consecutive days prior the test day by placing them on a treadmill (LE87 10RTS, Pan lab Harvard apparatus) at increasing speed at a 3 cm/sec rate. Mice were left to recover for 2 days before test day. For the test day, mice ran with a start speed of 8 cm/sec and every 2 min speed was increased at a 3 cm/sec rate. Belt slope was set at a 15° inclination, and the electric shock intensity of the resting pad was set at 0.2 mA. Mice ran until exhaustion signs were prominent, i.e. the presence of the mouse to the lower part of the lane, near the shock grid for more than 10 sec. Time, speed and distance ran were noted. For the grip strength test (76-1066, Harvard Apparatus), the mouse was lowered over the grid bar keeping the torso horizontal and allowing only its forepaws to attach to the sensor bar. The mouse was then gently pulled back by its tail ensuring it gripped the horizontal part of the bar, while the torso remained horizontal, and the maximal grip strength value was recorded. Each mouse was subjected to the same procedure at least 3 times per training day with a minimum 5-minute interval to allow the mice to recover. The grip test was repeated 2 days later, and the individual measurements were averaged per day per mouse.

### Statistical analysis

For comparison between two groups, unpaired student’s t-test was performed to calculate the p values. For comparisons between three groups One-way ANOVA was performed. P-values are indicated in each graph for the respective comparisons.

---

**Figure S1:**
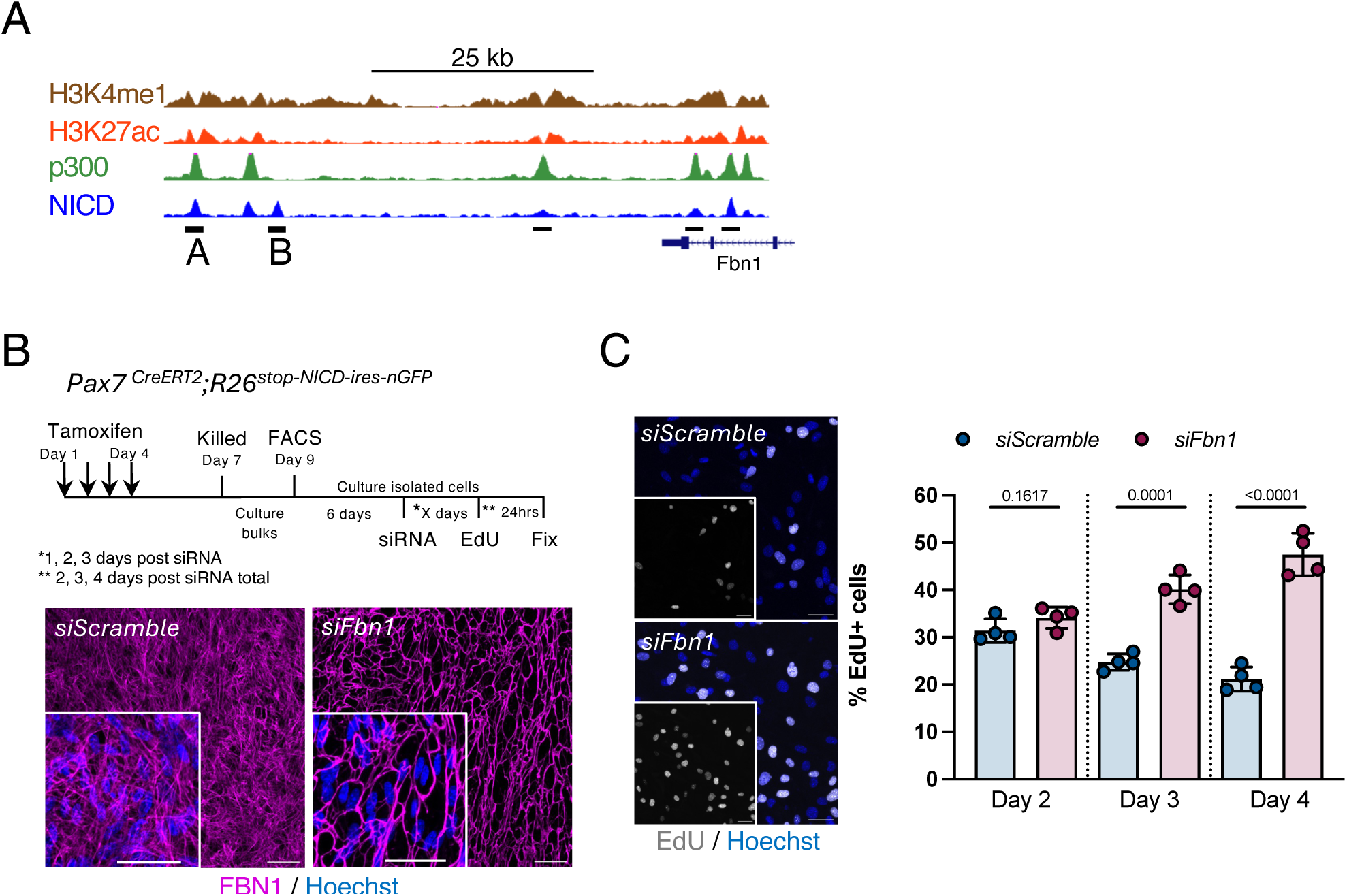
Transcriptional regulation of Fbn1 by Notch signaling. **A.** NICD-bound enhancers associated with mouse *Fbn1* loci co-occupied by the histone acetyltransferase p300 and also featuring the hallmark modifications of histone H3 Lys 4 monomethylation (H3K4me1) and/or histone H3 Lys 27 acetylation (H3K27ac); enhancers A and B were cloned, Fig. 1B). **B**. Experimental schematic for *Fbn1* downregulation in FACS-isolated satellite cells. Representative images of si*Scramble* and si*Fbn1* knock-down cultures 4 days post siRNA transfection. **C**. Quantification of EdU^+^ cells 2, 3, and 4 days post siRNA, and representative EdU immunofluorescence images at day 4 post siRNA. Unpaired *t-test* ±S.D.; scale bars = 40μm

**Figure S2:**
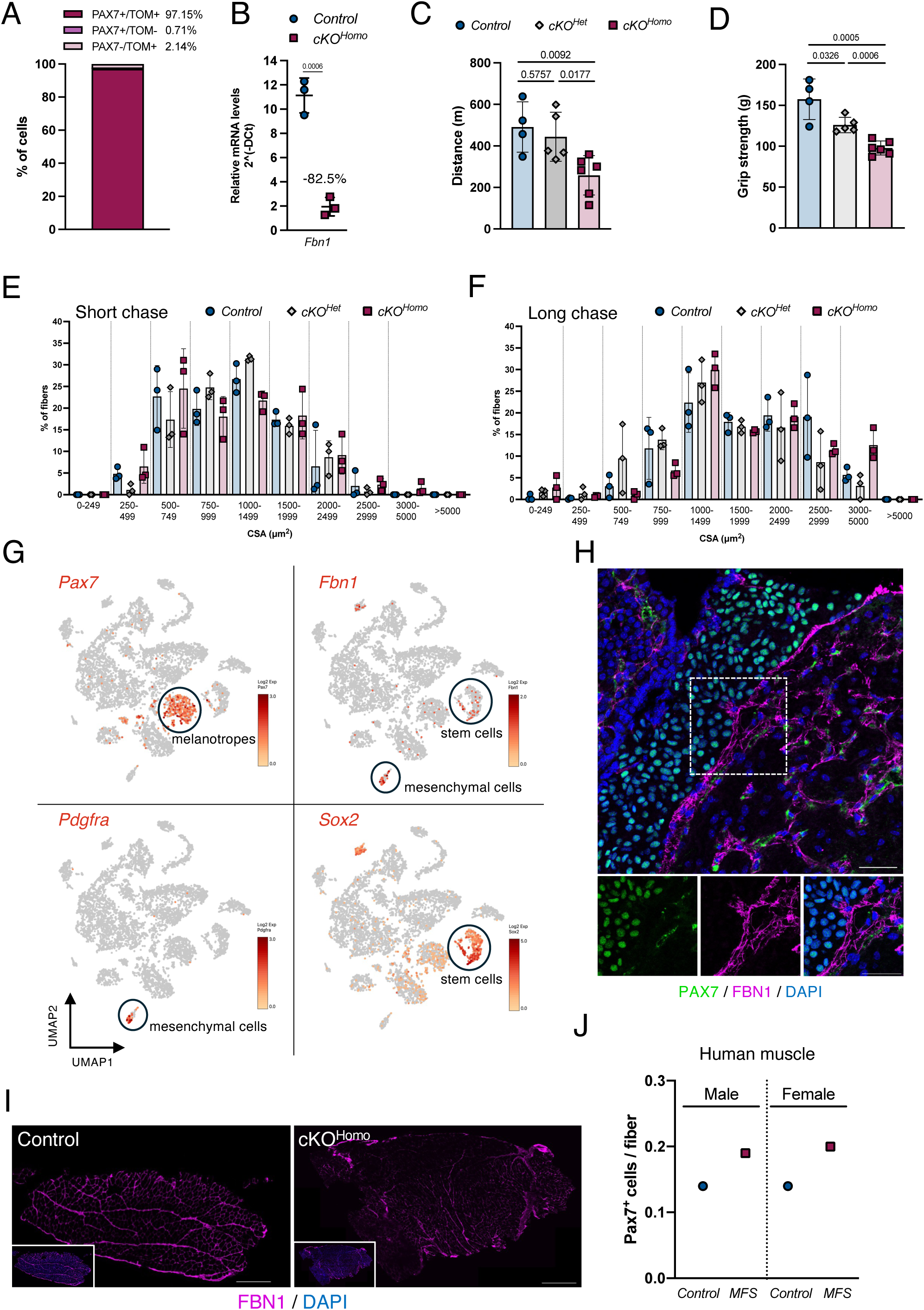
Characterization of Fbn1 deletion in Pax7^+^ muscle and pituitary gland cells. **A.** Percentage of cells expressing Pax7, Tom, or Pax7/Tom from control (*Pax7^CreERT2^;Fbn1^+/+^;R26^tdTom^*) mice (n = 3 mice) based on immunofluorescent staining of TA muscle sections (PAX7^+^/TOM^+^ vs PAX7^+^/TOM^-^ p=<0.0001; PAX7^+^/TOM^+^ vs PAX7^-^/TOM^+^ p=<0.0001; PAX7^+^/TOM^-^ vs PAX7^-^/TOM^+^ p=0.0322). **B.** RT-qPCR analysis of *Fbn1* from TOM^+^ isolated satellite cells from Control and cKO^Homo^ mice (n=3 mice/genotype) two weeks post tamoxifen treatment. **C.** Quantification of treadmill performance analysis in mice from the long-term Fbn1 depletion cohort (n = 4-6 mice/genotype). **D.** Quantification of grip strength performance from the long-term Fbn1 depletion cohort (n = 4-6 mice/genotype). **E.** Quantification of fiber cross-sectional area (CSA) from the short-chased mice (n = 3 mice/genotype). **F.** Quantification of CSA from the long-chased cohort (n = 3 mice/genotype). **G.** UMAP of single-cell RNAseq from pituitary gland cells indicating the expression of *Pax7*, *Fbn1*, *Pdgfra* and *Sox2* (Mayran et al., 2019). **H.** Immunostaining of pituitary gland sections from wild-type mice. PAX7^+^ cells in the intermediate lobe do not express FBN1 (magenta). **I.** FBN1 decrease in *Fbn1* cKO^Homo^ in FAPs (*Pdgfra^CreERT2^;Fbn1^fl/fl^;R26^tdTomato^*) compared to Control (*Pdgfra^CreERT2^;Fbn1^+/+^;R26^tdTomato^*) TA muscle sections (n = 3 mice/genotype; scale bar = 200 μm). **J.** Quantification of PAX7+ cells on human muscle biopsies. Unpaired *t-test* ±S.D (B); One-way ANOVA ±S.D. (C-F); scale bars = 40 μm.

**Figure S3:**
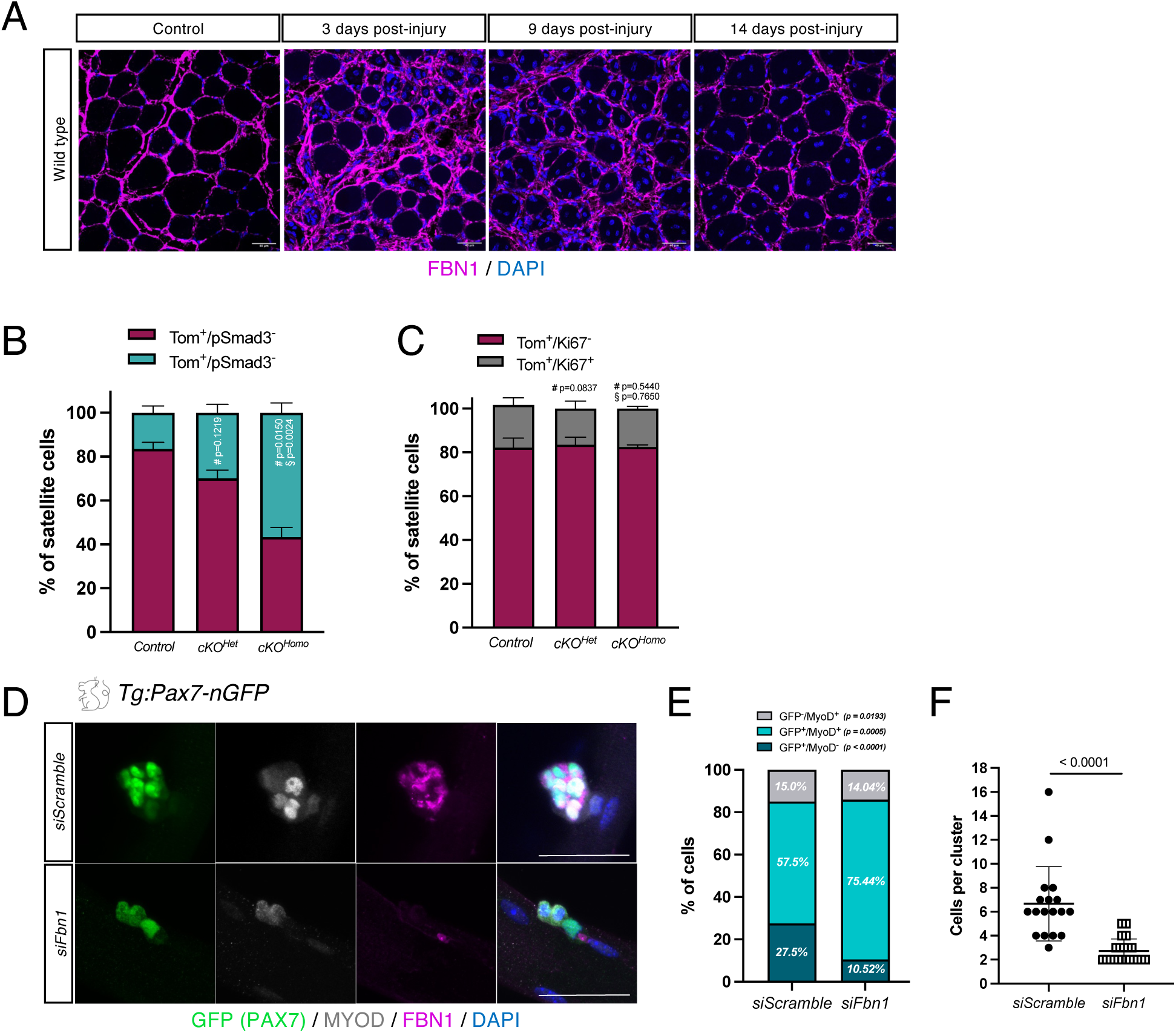
FBN1 knockdown impairs self-renewal in vitro and satellite cell recovery during regeneration. **A.** Immunostaining of TA muscle sections from wild-type mice at 0 days (control), 3, 9 and 14 days post TA injury. **B.** TOM^+^/pSMAD3^±^, and **C.** Tom^+^/Ki67^±^ satellite cells in control and *Fbn1* cKO mice (n = 3 mice per genotype) 14 days post injury. **D.** Immunostaining of 72hr-cultured individual myofibers from *Tg:Pax7-nGFP* mice transfected with *siScramble* (top panel) or siFbn1 (bottom panel). **E.** Quantification of GFP^+^/MyoD^-^, GFP^+^/MyoD^+^, and GFP^-^/MyoD^+^ cells from the resulted clusters (n = 3 per siRNA), and **F.** of cluster sizes. One-way ANOVA ±S.D.; # compared to Control; § compared to cKO^Het^ (B-C); unpaired *t-test* ±S.D (E-F); Scale bars = 40 μm.

**Figure S4:**
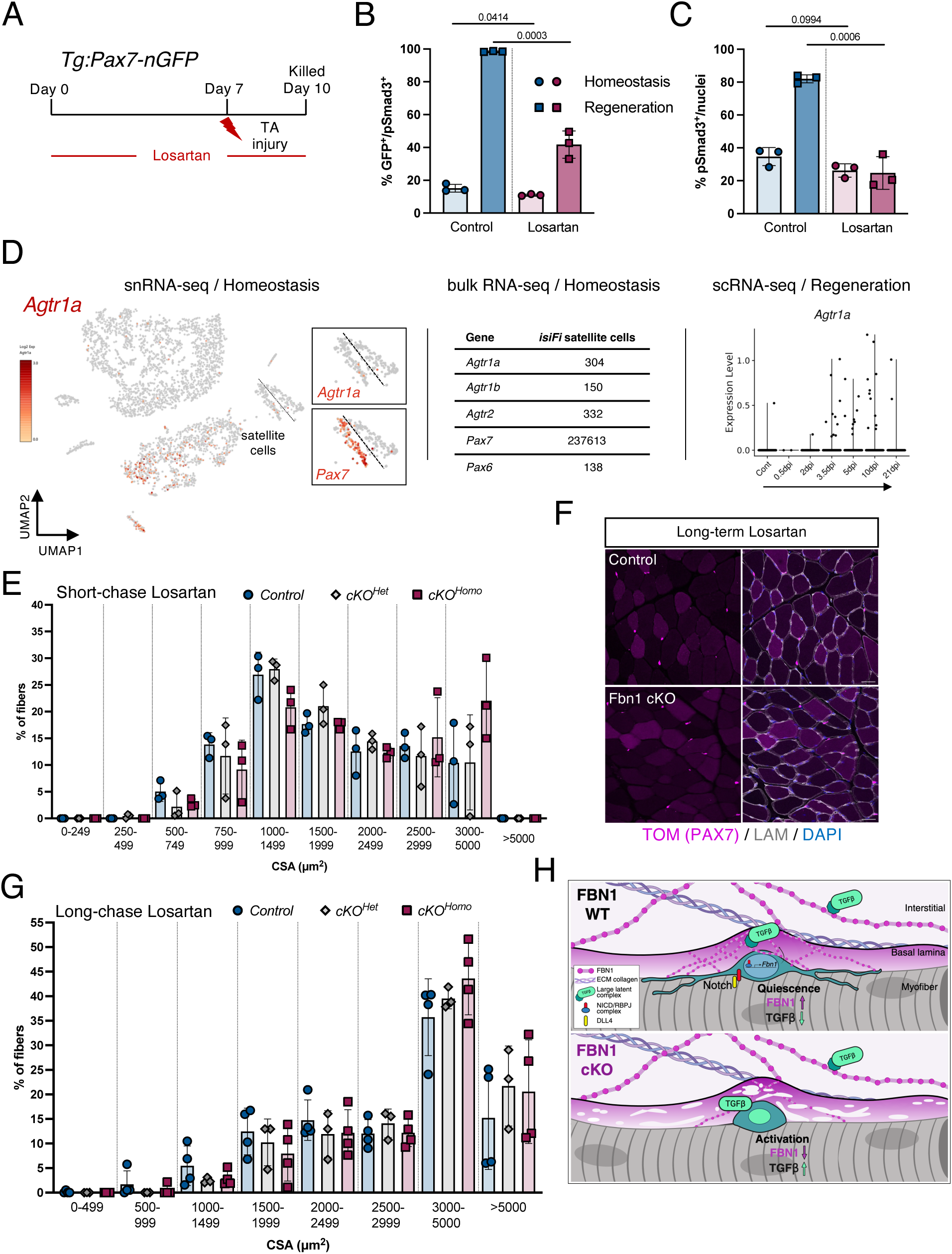
Angiotensin II type 1 receptor antagonist restores quiescence of Fbn1-deficient satellite cells and prevents muscle performance decline. **A.** Experimental scheme of losartan administration to *Tg:Pax7-nGFP* mice. **B.** Quantification of pSMAD3^+^ satellite cells (GFP^+^) and **C.** total pSMAD3^+^ nuclei (right graph). **D.** Expression of *Agtr1a* in snRNAseq from intact TA muscle (Machado et al., 2021), in bulk RNAseq on *in situ* fixed satellite cells from hind limb muscles (Machado et al., 2017) and scRNAseq from injured TA during regeneration (Oprescu et al., 2020). In the bulk RNAseq table, *Pax7* is shown as positive control and *Pax6*, which is not expresses in muscle, as background baseline of the reads). **E.** Quantification of fiber cross-sectional area (CSA) from the short-term. **F.** Immunostaining of TA muscle sections from Control and *Fbn1* cKO TA muscles from cohorts with and without losartan treatment (long-term exposure). and **G.** Quantification of fiber cross-sectional area (CSA) from the long-term losartan exposure in *Fbn1*-cKO cohorts (n = 3 mice per genotype). **H**. Proposed model of Notch-FBN1-TGFβ axis in the maintenance of skeletal muscle stem cell quiescence. Unpaired *t-test* ±S.D (B and D); one-way ANOVA ±S.D. (E and D); scale bars = 40μm.

## Notes

### Competing Interest Statement

The authors have declared no competing interest.

