## Supplemental Tables for "Muscle stem cells produce a protective Fibrillin-1 matrix to prevent precocious activation"

**Supplementary tables**

**Table S1**: List of primary antibodies.

| **Antibody** | **Reference** | **Dilution** |
|---|---|---|
| GFP chick | Abcam #13970 | 1/1000 |
| RFP rat | Chromotek #CR-5F8 | 1/250 |
| Fbn1 rabbit | Shi et al., 2021 | 1/1500 |
| Pax7 mouse | Santa Cruz #sc-81648 | 1/200 |
| MyoD mouse | Dako #M3512 | 1/200 |
| pSmad3 rabbit | Abcam #ab52903 | 1/200 |
| Ki67 rabbit | Abcam #ab16667 | 1/200 |
| Laminin conjugated | Biotechne #NB300-144AF647 | 1/200 |

**Table S2**: Coordinates of the *Fbn1*-associated enhancers (mouse) used in the luciferase assays.

| **Enhancer** | **Chromosome** | **Start-End** | **Size (bp)** |
|---|---|---|---|
| *Fbn1A* | Chr2 | 125,101,224-125,101,733 | 509 |
| *Fbn1B* | Chr2 | 125,105,576-125,106,100 | 524 |

**Table S3**: List of primers used for RT-qPCR (mouse).

| **Gene** | **Forward primer (5'-3')** | **Reverse primer (5'-3')** |
|---|---|---|
| *Fbn1* | TGGGTTCCAACTGGCATCAG | ATCCGTGTTCACACAGCGTC |

**Supplementary code S1**

**This code was implemented in R using the Seurat package to perform data loading, integration, preprocessing, visualization, and subclustering of single-cell RNA sequencing data.**

```
library(Seurat) #Version-5.3.0
library(dplyr) #Version-1.1.4
library(ggplot2) #Version-3.5.2

########LoadTheData###
base_dir <- "PATH"
# List of SRR project IDs
srr_list <- paste0("SRR102754", 13:19)
# List to store Seurat objects
seurat_objects <- list()
# Create Seurat objects for each SRR project
for (srr in srr_list) {
  data_dir <- file.path(base_dir, paste0(srr, "PATH"))
  mtx_file <- file.path(data_dir, "matrix.mtx.gz")
  barcodes_file <- file.path(data_dir, "barcodes.tsv.gz")
  features_file <- file.path(data_dir, "features.tsv.gz")
  count <- ReadMtx(mtx = mtx_file, cells = barcodes_file, features = features_file)
  seurat_object <- CreateSeuratObject(counts = count, project = srr)
  seurat_objects[[srr]] <- seurat_object
}
#
seurat_obj_cont <- seurat_objects[["SRR10275413"]]
seurat_obj_0.5dpi <- seurat_objects[["SRR10275414"]]
seurat_obj_2dpi <- seurat_objects[["SRR10275415"]]
seurat_obj_3.5dpi <- seurat_objects[["SRR10275416"]]
seurat_obj_5dpi <- seurat_objects[["SRR10275417"]]
seurat_obj_10dpi <- seurat_objects[["SRR10275418"]]
seurat_obj_21dpi <- seurat_objects[["SRR10275419"]]

#####Integration#####
seurat_list <- list(
  seurat_obj_cont,
  seurat_obj_0.5dpi,
  seurat_obj_2dpi,
  seurat_obj_3.5dpi,
  seurat_obj_5dpi,
  seurat_obj_10dpi,
  seurat_obj_21dpi
)

#####labeling#####
timepoints <- c("cont", "0.5dpi", "2dpi", "3.5dpi", "5dpi", "10dpi", "21dpi")
# Merge Seurat objects using merge function
merged_seurat <- merge(
  x = seurat_objects[[1]],
  y = seurat_objects[2:length(seurat_objects)],
  add.cell.ids = timepoints,
  project = "merged"
)
```

```
merged_seurat[["percent.mt"]] <- PercentageFeatureSet(merged_seurat, pattern = "^mt-")
merged_seurat <- subset(merged_seurat, subset = nFeature_RNA > 200 & nFeature_RNA < 8000 &
percent.mt < 5)
merged_seurat <- NormalizeData(merged_seurat)
merged_seurat <- FindVariableFeatures(merged_seurat)
merged_seurat <- ScaleData(merged_seurat)
merged_seurat <- RunPCA(merged_seurat)
merged_seurat <- RunUMAP(merged_seurat, dims = 1:30)
merged_seurat <- FindNeighbors(merged_seurat, dims = 1:10)
merged_seurat <- FindClusters(merged_seurat, resolution = 0.15)
DimPlot(merged_seurat, reduction = "umap", group.by = "seurat_clusters",label = TRUE) +
    ggtitle("UMAP Colored by Clusters")
  ggsave('Integrated_umap.png', width=15, height=15)
DimPlot(merged_seurat, reduction = "umap", group.by = "orig.ident") +
    ggtitle("Integrated UMAP")
DimPlot(merged_seurat, reduction = "umap", split.by = "orig.ident", ncol = 2) +
    ggtitle("UMAP Split by Timepoint")

####Targets-genes####
  genes_to_plot <- c("Pax7", "Fbn1", "Heyl", "Hey1", "Hes1", "Notch3",
"Myod1","Mki67","Calcr","Agtr1a","Agtr1b","Agtr1")
  plot_titles <- c("Pax7", "Fbn1", "HeyL", "Hey1", "Hes1", "Notch3",
"Myod1","Mki67","CalcR","Agtr1a","Agtr1b","Agtr1")
  display_names <- c("Pax7", "Fbn1", "HeyL", "Hey1", "Hes1", "Notch3",
"Myod1","Mki67","CalcR","Agtr1a","Agtr1b","Agtr1")
  # re-order
  ordered_labels <- c("Cont", "0.5dpi", "2dpi", "3.5dpi", "5dpi", "10dpi", "21dpi")
  # Vlnplot-for-each
  for (i in seq_along(genes_to_plot)) {
    gene <- genes_to_plot[i]
    disp <- display_names[i]
    expr_data <- FetchData(merged_seurat, vars = gene, slot = "data")
    cells_keep <- rownames(expr_data)[expr_data[[gene]] > 0]
    # subset
    tmp <- subset(merged_seurat, cells = cells_keep)
    # Vlnplot
    p <- VlnPlot(
      tmp,
      features = gene,
      group.by = "orig.ident",
      slot = "data",
      pt.size = 0.5
    ) +
      scale_x_discrete(limits = ordered_labels) +
      ggtitle(bquote(italic(.(disp)))) +
      theme(
        axis.text.x    = element_text(angle = 45, hjust = 1),
        plot.title     = element_text(size = 14, face = "italic", hjust = 0.5),
        plot.margin    = margin(5, 5, 5, 5), legend.position = 'none'
      )
    # Save
    file_name <- paste0("Vln_", gene, ".png")
    ggsave(
      filename = file_name,
```

```
        plot      = p,
        width     = 5,
        height    = 5,
        dpi       = 300
    )
  }

#####Subclustering#####
  seurat_subset <- subset(merged_seurat, idents = c("Pax7-enriched-cluster"))   # select cluster
  seurat_subset <- NormalizeData(seurat_subset)
  seurat_subset <- FindVariableFeatures(seurat_subset, selection.method = "vst", nfeatures = 2000)
  seurat_subset <- ScaleData(seurat_subset, features = rownames(seurat_subset))
  seurat_subset <- RunPCA(seurat_subset)
  seurat_subset <- FindNeighbors(seurat_subset, dims = 1:10)
  seurat_subset <- FindClusters(seurat_subset, resolution = 0.25)
  seurat_subset <- RunUMAP(seurat_subset, dims = 1:10)

  # Integrated MuSC-UMAP_orig.ident
  p1 <- DimPlot(seurat_subset, reduction = "umap", group.by = "orig.ident") +
    ggtitle("Integrated MuSC-UMAP")
  print(p1)
  ggsave("Integrated MuSC-UMAP_orig.ident.png", plot = p1, width = 6, height = 6, dpi = 300)
  # MuSC-UMAP Colored by Clusters
  p2 <- DimPlot(seurat_subset, reduction = "umap", group.by = "seurat_clusters", label = TRUE) +
    ggtitle("MuSC-UMAP Colored by Clusters")
  print(p2)
  ggsave("MuSC-UMAP Colored by Clusters_by_clusters.png", plot = p2, width = 6, height = 6,
dpi = 300)
  VlnPlot(seurat_subset, features =
c("Pax7","Hs6st3","Calcr","Myod1","Myf5","Myog","Ctnnb1","Zbtb18","Mycl","Scx","Mef2a","H
es6", "E2f8","Klf5","Id1","Tead4","Piezo1","Piezo2","Mymx","Acta1","Cdkn1c","Myh1","Acta1"))
  ggsave('MuSC-myogenic_vlnplot.png', width=15, height=15)
  FeaturePlot(seurat_subset, features =
c("Pax7","Hs6st3","Calcr","Myod1","Myf5","Myog","Ctnnb1","Zbtb18","Mycl","Scx","Mef2a","H
es6","E2f8","Klf5","Id1","Tead4","Piezo1","Piezo2","Mymx","Acta1","Cdkn1c","Myh1","Acta1"))
  ggsave('MuSC-myogenic_umap.png', width=15, height=15)

#####Subclustering-MuSC-> qMuSCandactMuSC#####
  qMuSC <- subset(seurat_subset, idents = c(0, 1, 2, 5))
  actMuSC <- subset(seurat_subset, idents = c(3, 4))
  genes_to_plot <- c("Pax7", "Fbn1", "Heyl", "Hey1", "Hes1", "Notch3",
"Myod1","Mki67","Calcr","Agtr1a")
  display_names <- c("Pax7", "Fbn1", "HeyL", "Hey1", "Hes1", "Notch3",
"Myod1","Mki67","CalcR","Agtr1a")
  ordered_labels <- c("Cont", "0.5dpi", "2dpi", "3.5dpi", "5dpi", "10dpi", "21dpi")
  object_list <- list(qMuSC = qMuSC, actMuSC = actMuSC)
  #
  for (obj_name in names(object_list)) {
    obj <- object_list[[obj_name]]
    for (i in seq_along(genes_to_plot)) {
      gene <- genes_to_plot[i]
      disp <- display_names[i]
      # VlnPlot
      p <- VlnPlot(
```

```
        obj,
        features = gene,
        group.by = "orig.ident",
        slot = "data",
        pt.size = 1
    ) +
        scale_x_discrete(limits = ordered_labels) +
        ggtitle(bquote(italic(.(disp)))) +
        theme(
            axis.text.x = element_text(angle = 45, hjust = 1, family= "Helvetica"),
            plot.title = element_text(size = 18,face = "bold.italic", hjust = 0.5, family= "Helvetica"),
            plot.margin = margin(5, 5, 5, 5),
            legend.position = 'none'
        )
    # Save
    file_name <- paste0(obj_name, "-Vln_", gene, ".png")
    ggsave(
        filename = file_name,
        plot = p,
        width = 4,
        height = 4,
        dpi = 300
    )
  }
}
```
